# Refining fine-mapping: effect sizes and regional heritability

**DOI:** 10.1101/318618

**Authors:** Christian Benner, Aki S. Havulinna, Veikko Salomaa, Samuli Ripatti, Matti Pirinen

## Abstract

Recent statistical approaches have shown that the set of all available genetic variants explains considerably more phenotypic variance of complex traits and diseases than the individual variants that are robustly associated with these phenotypes. However, rapidly increasing sample sizes constantly improve detection and prioritization of individual variants driving the associations between genomic regions and phenotypes. Therefore, it is useful to routinely estimate how much phenotypic variance the detected variants explain for each region by taking into account the correlation structure of variants and the uncertainty in their causal status. Here we extend the FINEMAP software to estimate the effect sizes and regional heritability under the probabilistic model that assumes a handful of causal variants per each region. Using the UK Biobank data to simulate GWAS regions with only a few causal variants, we demonstrate that FINEMAP provides higher precision and enables more detailed decomposition of regional heritability into individual variants than the variance component model implemented in BOLT or the fixed-effect model implemented in HESS. Using data from 51 serum biomarkers and four lipid traits from the FINRISK study, we estimate that FINEMAP captures on average 24% more regional heritability than the variant with the lowest P-value alone and 20% less than BOLT. Our simulations suggest how an upward bias of BOLT and a downward bias of FINEMAP could together explain the observed difference between the methods. We conclude that FINEMAP enables computationally efficient estimation of effect sizes and regional heritability in the era of biobank scale data.

## Introduction

Over the last decade, Genome-Wide Association Studies (GWAS) have accumulated a massive catalog of robust statistical associations between numerous genomic regions and complex traits or diseases^1^. Typically, genetic variants highlighted by GWAS explain only a small fraction of total phenotypic variation (the missing heritability problem^2^) and are clustered into clumps of even hundreds of highly correlated variants giving rise to the fine-mapping problem^3^.

The missing heritability problem has spurred highly successful methodological research in variance components models to evaluate the heritability contribution from the whole genome, not just from the statistically significantly associated variants^4-6^. Software packages implementing variance component models include EMMAX^7^, GCTA^8^, GEMMA^9^, MMM^10^ and BOLT^5^. Common to these implementations is the requirement of individual-level genotype-phenotype data and the assumption that the phenotype has a polygenic architecture under which all variants are causal with tiny effects. Heritability estimation using a polygenic model has also been carried out with LD Score regression^6^ and a polygenic score method AVENGEME^11^ from GWAS summary statistics without access to the original genotype-phenotype data. Together these methods have considerably narrowed the gap between estimated SNP heritability and heritability estimates from twin studies for many phenotypes and, importantly, shed light on the genetic architecture and enrichment of functional genetic variation in complex traits and diseases^12,13^.

With increasing statistical power of GWAS comes a possibility to narrow the heritability down into specific regions of the genome. Recently, heritability estimation in genomic regions from GWAS summary statistics was introduced in the software package HESS^14^ under the assumption of arbitrary genetic architecture and a fixed-effect model for causal effect sizes. HESS estimates heritability through a regularized quadratic form built on marginal effect size estimates of all variants and their pairwise correlations to account for Linkage Disequilibrium (LD) between them. HESS was shown^14^ to be more accurate than LD Score regression in regional heritability estimation on simulated data when both methods were run with LD information either from the original genotype data or a reference panel.

Although polygenic and fixed-effect models have been very useful approaches for the estimation of genome-wide or regional heritability, the model assumptions of polygenicity or arbitrary genetic architecture likely leave room for improvement in accuracy if regional heritability can be attributed to only a few causal variants. Furthermore, our ultimate goal is to pinpoint the individual variants contributing to heritability and obtain accurate effect size estimates for them. Because existing heritability estimation methods do not provide such information, we turn to the fine-mapping framework instead.

The decisive solution to the fine-mapping problem is often hindered by the lack of information due to small effect sizes, high LD between variants and imperfect genotype information. Therefore, many recent fine-mapping methods^15-19^ implement principled probabilistic quantification of causal variants that can then be used for downstream analyses. As GWAS sample sizes are soon counted in millions providing unprecedented accuracy for statistical fine-mapping, it would be useful to routinely evaluate how much of the regional heritability can be explained by the fine-mapped variants. In particular, we may expect that fine-mapping captures a large part of the regional heritability in 1) molecular endophenotypes with moderate to large effect sizes at causal variants, and 2) in large-scale biobank projects and ongoing GWAS meta-analyses with adequate statistical power to detect most individual causal variants even with smaller effect sizes. In cases where the fine-mapping seems to explain the total regional heritability, we have a much more informative picture of the regional architecture than is given by the existing heritability estimation methods that do not identify individual causal variants. Therefore, we have extended the FINEMAP software to estimate the effect sizes and regional heritability by accounting for the uncertainty of the causal configuration and the LD-adjusted joint effect sizes of causal variants.

We compare regional heritability estimation using FINEMAP with both the variance component model implemented in BOLT and the fixed-effect model implemented in HESS. Ideally, with such comparisons, we can evaluate how much of the total regional heritability we capture with fine-mapped variants. Previously, Gusev et al.^20^ have studied how the variance explained by the variant with the lowest P-value of a GWAS region compares to the results of a variance component model applied to the same region. They reported that on average the variance component model explained 1.29 times more heritability over nine diseases. We use 51 serum biomarkers and four lipid traits from the FINRISK study, with a broad spectrum of genome-wide heritability levels, both for regional heritability estimation and to report fine-mapping results of these phenotypes at 109 genomic regions. As an example where both FINEMAP and BOLT give a similar estimate of about 70% regional heritability, we study in detail the association between the *PON1* gene (MIM: 168820) and Paraoxonase 1 levels. To interpret the results from our biomarker data analysis more generally, we also perform a simulation study to evaluate the methods in genomic regions that contain only a handful of causal variants since BOLT and HESS have been previously evaluated mainly on highly polygenic architectures^14,21^. As our ultimate goal is to narrow down the polygenic regional heritability into individual variants, it is useful to know how the heritability estimates of FINEMAP, BOLT and HESS relate to each other in cases where such decomposition is indeed possible. More generally, an ability to do routine comparisons between the regional heritability estimates originating from different modeling assumptions, including a fine-mapping model, further informs us about the genetic architecture of each region.

## Materials and Methods

### FINEMAP model

The goal of statistical fine-mapping is to refine a large set of genetic variants simply associated with a phenotype down to a much smaller set of variants with direct effect on the phenotype by accounting for the complex correlations between the variants. Although we call the variants with such direct effects “causal”, we bear in mind that further laboratory experiments would be required to establish the exact biological mechanisms of those variants.

The statistical framework in FINEMAP (see Web Resources) to infer causal variants is built on Bayesian variable selection for linear regression models^22^ using Bayes factors. Previously^18^, we derived the Bayes factor to assess the support for each combination of *k* variants denoted by *γ*, and hereafter called causal configuration, over the null configuration. Our previous derivation started with a likelihood approximation for the joint linear regression model for all variants in the region and assumed small enough effect sizes that we could ignore the differences in residual variance between causal configurations. In the following, we derive a corresponding Bayes factor by using a likelihood approximation of only the causal variants, which makes the effect size estimation feasible, and by allowing different residual variances for different configurations. For this, we assume the following Bayesian linear regression model for the causal configuration *γ*,

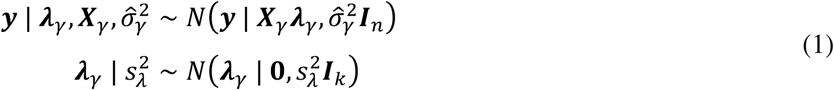

where ***y*** is a standardized vector with phenotypic values for *n* individuals (with Ê[***y***] = 0, 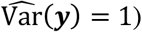, ***X***_*γ*_ is a matrix of dimension *n*× *k* with standardized genotypes at the variants, ***λ***_*γ*_ are the additive effect sizes of the standardized variants, 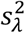 is the user-given prior variance for the effect sizes and 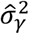, is the point estimate of the residual variance in the multiple linear regression that we learn from the data (details below). Integrating out effect sizes ***λ***_*γ*_ analytically yields the marginal likelihood of the causal configuration

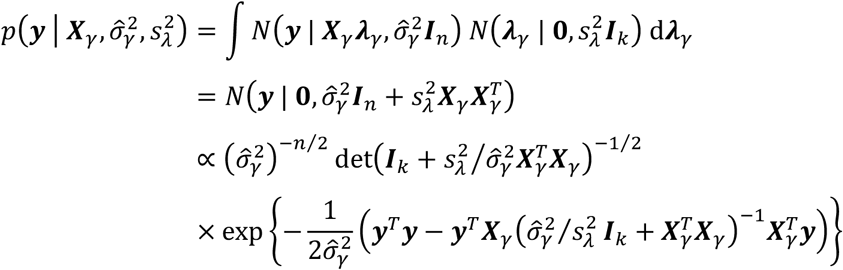

where we reduced computational complexity by applying the Matrix determinant lemma and Woodbury matrix identity in line 2. The computational complexity of evaluating the marginal likelihood can be further reduced substantially by replacing the phenotype-genotype data with GWAS summary statistics and pairwise correlations between variants^15-19^. That is, we replace individual-level data (***y***, ***X***) with GWAS summary data (***ẑ***, ***R̂***)whence

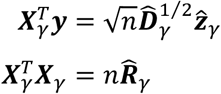

where ***ẑ***_*γ*_ is a vector of ratios of effect size estimates and their standard errors from individual univariate linear regressions of each variant in the causal configuration, ***R̂***_*γ*_ is a matrix of pairwise sample correlations among the variants and ***D̂*** is a diagonal matrix of estimates for residual variances in the univariate linear regressions. Thus, we can compute the Bayes factor to assess the support for a causal configuration *γ* over the null configuration *γ*_0_, that contains no variants, by using only GWAS summary data on variants included in the causal configuration *γ*

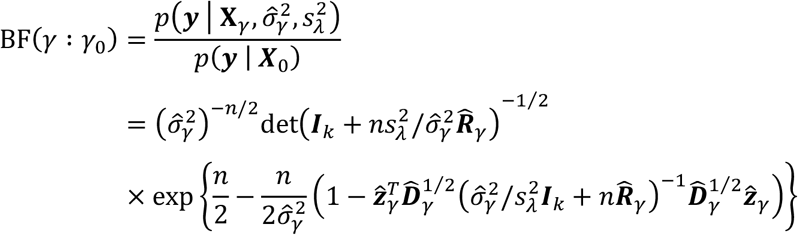

Numerically, this Bayes factor agrees with our earlier derivation^18^ when causal variants have small effect sizes and hence 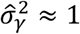. With the current derivation of the Bayes factor, we can also account for causal variants with larger effect sizes by allowing for different residual variances for different configurations and provide a simple way to extract the posterior distribution of the effect sizes as explained next.

### Estimation of effect sizes and regional heritability

The posterior distribution of the effect sizes of variants included in a causal configuration is

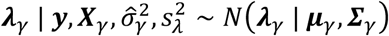

Where 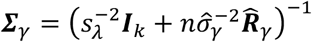 and 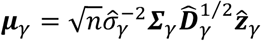. An estimate of the heritability of a causal configuration *γ* is given by the ratio between the variance explained by the causal variants and the total variance of the phenotype, which we here compute conditional on the effect sizes ***λ***_*γ*_ as

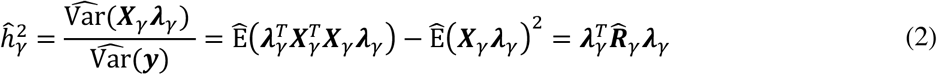

where the empirical estimate from the data 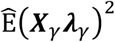 equals zero because of centered genotypes. Hence our point estimate is 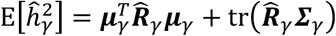 and we compute the posterior distribution of 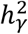, by evaluating Equation (2) for each draw from the posterior for ***λ***_*γ*_. This approach allows us to obtain the mean, standard deviation and 95% credible interval (equal tail area interval) of the posterior. We expect high quality posterior inference if ***R̂***_*γ*_ is a good approximation to the pairwise sample correlations computed from the original GWAS genotype data^23^.

### Large effect sizes

For quantitative traits, keeping estimates for residual variances in ***D̂*** fixed to one is a good approximation in genomic regions with small causal effects and FINEMAP v1.1 uses this approximation. However, this approach can lead to problems when causal variants have larger effect sizes^24^. In FINEMAP v1.2, we account for different magnitudes of effect sizes by calculating, for each variant *ℓ*, the maximum likelihood estimate

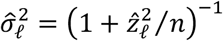

for the residual variance in univariate linear regression. If all variants individually account for less than heritability of 1%, we keep 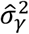, in Equation (1) fixed at a value of 1.0 and set the prior variance for the effect size 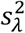 equal to 0.05^2^; this means that a causal variant contributes less than heritability of 1% with probability 0.95^18^. Otherwise, we expect to see larger effect sizes. In this case, we set 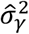 in Equation (1) equal to the maximum likelihood estimate for the residual variance from multiple linear regression

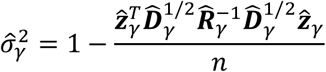

of variants in the causal configuration *γ*. For each causal configuration, we further use an equidistant grid of four values for the prior variance 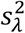 with a lower bound at 0.05, and an upper bound where, with probability 0.95, a causal variant contributes less than the point estimate of heritability 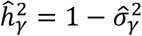 of a causal configuration *γ* of the variants with pairwise absolute correlations less than 0.95 that are marginally the most significant and that each contribute more than heritability of 1%. For each grid value, we compute the Bayes factor for assessing the evidence against the null configuration. We obtain the final Bayes factor by computing the average value of the individual Bayes factors. This model averaging corresponds to a uniform prior distribution on 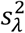 over the grid points.

Our strategy for adapting to large effect sizes requires no actions from the user and therefore differs from the approach^24^ of CAVIARBF v0.2.1 (see Web Resources) which requires user-given values for the prior standard deviation *s_λ_*. We compare FINEMAP v1.2 with CAVIARBF and FINEMAP v1.1 on simulated datasets with three causal variants among 257 variants in a genomic region (see below “GWAS regions in UKBB data”). Each dataset was analysed with FINEMAP and CAVIARBF using default values and allowing for a maximum of four causal variants in the region. For CAVIARBF, we excluded the null configuration from the output and used multiple values 0.05, 0.1, 0.2, 0.4, 0.8 to specify the prior standard deviation *s_λ_* as recommended^24^.

### Bayesian model-averaging

The Bayesian variable selection approach to fine-mapping causal variants is a hard combinatorial problem which precludes exhaustive enumeration of all possible causal configurations. In previous work^18^, we tackled the combinatorial problem with an ultra-fast Shotgun Stochastic Search^25^ algorithm to identify a list of causal configurations which captures a large majority of the total posterior probability^18^. According to the Bayesian paradigm, we account for model uncertainty through Bayesian Model Averaging (BMA)^26^. BMA combines inference about a quantity common to all models by averaging the posterior distribution of that quantity under each model using posterior model probabilities as weights.

Using BMA, we compute the marginal posterior inclusion probability (PIP) for the *ℓ*th variant by summing up the posterior probability of all causal configurations in which this variant is included. We model-average estimates of heritability to quantify the contribution of causal variants. Examples of other useful model-averaged estimates are the posterior effect size mean and standard deviation of the *ℓ*th variant. We output marginalized shrinkage estimates of the posterior effect size mean and standard deviation by averaging over all causal configurations assuming that the effect size of the *ℓ*th variant is zero if the variant is absent from a causal configuration. We also output the conditional estimates of the posterior effect size mean and standard deviation conditional on inclusion by averaging over only those causal configurations in which the *ℓ*th variant is included. The magnitude of the conditional model-averaged estimate of the *ℓ*th variant is much larger than the shrinkage model-averaged estimate if the PIP for the *ℓ*th variant is small.

### Polygenic and fixed-effect models

Variance component models are extensively used for heritability estimation under a polygenic model assumption. We performed regional heritability estimation from individual-level genotype-phenotype data with BOLT v2.3 (see Web Resources) using default settings. In genome-wide analyses, we used an LD pruned set of 834,334 variants obtained after two rounds of pruning at *r*^2^ = 0.5 in PLINK (--indep-pairwise 100 5 0.5, see Web Resources) while in regional analyses we used an unpruned set of variants and LD stratification^12^ by assigning variants to different variance components based on quartiles of LD scores to account for heterogeneity in LD and Minor Allele Frequency (MAF).

Since the model assumption of polygenicity may not hold in any particular genomic region^14^, we also estimated regional heritability with HESS v0.5.2 (see Web Resources) to avoid assumptions about causal effect sizes. HESS requires GWAS summary statistics and the pairwise correlations between the variants to account for LD. We ran HESS with the default settings using LD information obtained from the original GWAS and command-line flag --min-maf 0.01 for MAF filtering. We also ran HESS without regularization of the LD matrix by additionally setting the command-line flags --max-num-eig *m* and --min-eigval 0.0 to use the full Singular Value Decomposition SVD of the LD matrix, where *m* is the number of variants in the genomic region.

### Cohorts

We used data from the FINRISK27 study and UK Biobank^28^ (UKBB). FINRISK is a representative, cross-sectional survey of the Finnish working age population. Since 1972, a random sample of 6,000-8,000 individuals has been collected every five years to study risk factors of chronic diseases. The study protocols of the FINRISK study surveys used in this work (1992, 1997, 2002, 2007, 2012) were approved by the Ethics Committee of the National Public Health Institute until 1997 and after that by the Ethics Committee of Helsinki and Uusimaa Hospital District. UKBB is a longitudinal study of individuals between 40 to 69 years of age in the United Kingdom approved by the North West Multi-centre Research Ethics Committee. From 2006 to 2010, a sample of 500,000 individuals was collected to investigate genetic and environmental factors involved in disease development. All participants of the FINRISK study and UKBB have given an informed consent.

### Genotype data

We used genotype data on 1) 27,294 individuals included in FINRISK 1992-2012, and 2) 408,966 “white British” individuals provided by UKBB. For FINRISK, imputation to 15 million variants was performed with IMPUTE2 (see Web Resources) using a Finnish imputation panel from high-pass whole-genome sequence data. For documentation about imputation of the UKBB genotype data see Web Resources. In both datasets, we removed variants with MAF less than 1% and excluded related individuals with KING’s^29^ kinship coefficient greater than 0.0442 using a greedy algorithm that maximizes the final sample size. This left us with 22,640 samples in FINRISK and 343,740 samples in UKBB. In FINRISK data, we retained biallelic variants with less than 5% missing calls and imputation quality greater than 0.9. In the UKBB data, we retained genotyped biallelic variants with imputation quality equal to 1.0 that are part of the Haplotype Reference Consortium panel (see Web Resources).

### Phenotype data

In our analyses, we used four common lipid measurements: low-density lipoprotein cholesterol (LDL), high-density lipoprotein cholesterol (HDL), total cholesterol (TC) and triglycerides (TG) on 21,320 unrelated individuals included in FINRISK 1992-2012 and serum levels of 51 biomarkers representing inflammation, metabolism, vascular function, oxidative stress, coagulation, renal function, angiogenesis, and myocardial necrosis from 5,267 unrelated individuals included in the FINRISK 1997 (Table S1). Details about lipid and biomarker measurements are given in ^30^ and ^31^. Individuals on lipid-lowering medication were excluded from all analyses. Each phenotype was adjusted by sex, age, age^2 and the first 15 principal components of genetic population structure using a linear regression model. Principal component analysis was performed separately for FINRISK 1992-2012 and FINRISK 1997 from the estimated genetic relatedness matrix computed from a set of LD pruned variants outside long-range LD regions with MAF greater than 5% and imputation quality score above 0.95. Regression residuals were normalized using a rank-based transformation to the standard Normal distribution.

### Biomarker and lipid-associated regions in FINRISK data

We used 5,267 individuals included in FINRISK 1997 and a linear model implemented in SNPTEST2 (see Web Resources) to test for genome-wide associations with 51 serum biomarkers and four lipid traits. We defined trait-associated genomic regions by first centering a 2 Mb window around the variant with the lowest P-value and then iteratively around the variant with the next lowest P-value not yet included in any chosen region until no further variant reached the genome-wide significance threshold of 5 × 10^−8^. Final genomic regions were obtained by merging physically overlapping regions. In the Paraoxonase 1 associated region, we performed a stepwise conditional analysis implemented in SNPTEST2 by first conditioning on the variant with the lowest P-value, hereafter called the lead variant, and then iteratively adding to the model the variant with the lowest conditional P-value until no further variant reached the genome-wide significance threshold or until we had conducted a pre-specified number of iterations. We estimated regional heritability with BOLT, HESS and FINEMAP. For FINEMAP we allowed for ten causal variants. We further ran FINEMAP allowing for one causal variant to estimate the heritability captured by the lead variant. To evaluate the effect of samples size on the regional heritability captured by fine-mapped variants, we estimated regional heritability with BOLT, HESS and FINEMAP for the four lipid traits both using 5,114 individuals included in FINRISK 1997 and 21,320 individuals in FINRISK 1992-2012.

### GWAS regions in UKBB data

To assess how different methods estimate regional heritability in a case where there are only a few causal variants in the region, we performed comprehensive simulations over genomic regions chosen from GWAS meta-analyses for coronary artery disease (CAD)^32^, Crohn’s disease (CD)^33^, lipid traits (LIPs)^34^, schizophrenia (SCZ)^35^, and type 2 diabetes (T2D)^36^. For each study, we retained the lead variants outside the human leukocyte antigen (HLA) region with a marginal P-value less than 5 × 10^−8^ and selected 100 lead variants (18 from CAD, 20 from CD, 21 from LIPs, 21 from SCZ, and 20 from T2D) for further analyses. Around each lead variant, we defined genomic regions of 3 Mb comprising variants with pairwise absolute correlations less than 0.95. For 98 regions with at least 200 variants in UKBB data, we considered the following fine-mapping setting:

- **Scenario H** Increasing regional heritability (*h^2^* = 0.0015, 0.05, 0.5)
- **Scenario N** Increasing GWAS sample size (*n* = 5,000, 50,000, 300,000)

A total of 980 datasets (10 replicates per each region) were generated under each of the 9 combinations of scenarios H and N using the following linear model

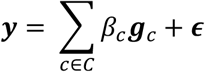

where *C* is the set of causal variants, ***g***_*c*_ the vector of standardized genotypes from 343,740 UKBB individuals at the *c*th causal variant, *β*_*c*_ the standardized effect size of the *c*th variant and ***ϵ*** is Normal noise with mean 0 and variance 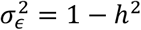.

The number of causal variants was three in each dataset. Causal variants were randomly chosen and their joint effect sizes were specified so that in the true causal configuration the three variants accounted for 61.8%, 25.8% and 12.4% of the regional heritability, respectively. These per-variant proportions were chosen from a meta-analysis of the association between *MC4R* locus (MIM: 155541) and body mass index^37^.

For each set of 980 simulations, we report the average heritability estimates. We assess the uncertainty quantifications of heritability estimates by comparing the standard deviation of heritability estimates with the average standard error (BOLT, HESS) or posterior standard deviation (FINEMAP). For 95% credible and confidence intervals (normal approximations), we calculate their coverage as the proportion of intervals that covered the true level of heritability. We also compute the root mean square error (RMSE) defined as the square root of the average squared deviation of the estimated heritability from the true level of heritability. RMSE summarizes both bias and precision of the estimation procedures with smaller RMSE values indicating better performance.

### Results

We start with a motivating example of a region with a large contribution to genome-wide heritability. Next, we study by simulation how different methods perform more generally. Finally, we apply the methods to the biomarker dataset.

### Regional heritability of paraoxonase 1 in FINRISK data

Paraoxonase 1 (PON1) is an HDL-associated enzyme, encoded by the *PON1* gene (MIM: 168820) on chromosome 7, whose serum levels have been reported to be reversely associated with systemic oxidative stress and atherosclerosis risk^38^. Using data on 4,613 individuals from FINRISK 1997, BOLT reported a high level of genome-wide heritability (*h*^2^ = 1.00, CI_0.95_: 0.83-1.00) in the PON1 measurements available in our biomarker panel. The only genome-wide significant (P < 5 × 10^−8^) variants were located near the *PON1* gene, which agrees with an earlier GWAS^38^. Therefore, we applied FINEMAP on a 7.5Mb genomic region around the *PON1* gene with 16,189 variants. FINEMAP estimated that with over 95% probability 5 variants are needed to explain the association signal and the fine-mapped variants together explain a heritability of *h*^2^ = 0.69 (CI_0.95_: 0.67-0.72) which is considerably more than the lead variant rs1157745 alone (*h*^2^ = 0.58, CI_0.95_: 0.56-0.61). The top configuration from FINEMAP included 5 variants (rs3917550, rs1157745, rs854559, rs2299260, rs705379) which we compared to the top 5 variants identified by the stepwise conditioning. The maximized likelihood was 54 times higher for the configuration identified by FINEMAP demonstrating the efficiency of the search algorithm of FINEMAP compared to the stepwise conditional analysis. BOLT estimated regional *h*^2^ = 0.70 (CI_0.95_: 0.66-0.74) which was similar to the estimate from FINEMAP but with a wider confidence interval. HESS with the default regularization gave an unexpectedly small value *h*^2^ = 0.51, that was clearly smaller than the marginal heritability contribution of the lead variant alone, with an extremely large standard error (SE = 6833.9). HESS without regularization of the LD matrix yielded an infeasible value *h*^2^ = 7.83 with a negative standard error.

As a conclusion, the *PON1* region is an example of a large effect region where FINEMAP provides fine-mapping that is statistically preferred over both the stepwise conditioning and single-variant analysis, and produces heritability estimates that are more detailed and more precise than the results from BOLT or HESS. However, since the *PON1* region is an extreme case with a very large regional heritability contribution, we next performed simulation studies to assess more generally the differences between FINEMAP, BOLT and HESS in regions with only a few causal variants.

### Simulations with GWAS regions in UKBB data

Using the datasets on 98 GWAS regions, we evaluated heritability estimation implemented in FINEMAP, HESS and BOLT in simulations with three randomly chosen causal variants. This setting is particularly suitable for FINEMAP and we are interested in how HESS and BOLT perform relative to FINEMAP in order to later interpret our biomarker data results.

The FINEMAP results verified our implementation as the estimates were nearly unbiased and the precision increased with the sample size (Figure 1 and Table S2A). For the smallest sample size of *n* = 5,000, FINEMAP underestimated the heritability for *h*^2^ of 5% and 50% but this effect disappeared with increasing sample size. This phenomenon shows how our prior distributions favor sparse models and smaller effect sizes in cases where the information in the data is weak, but how the effect of the priors diminishes with more informative data.

**Figure 1.**
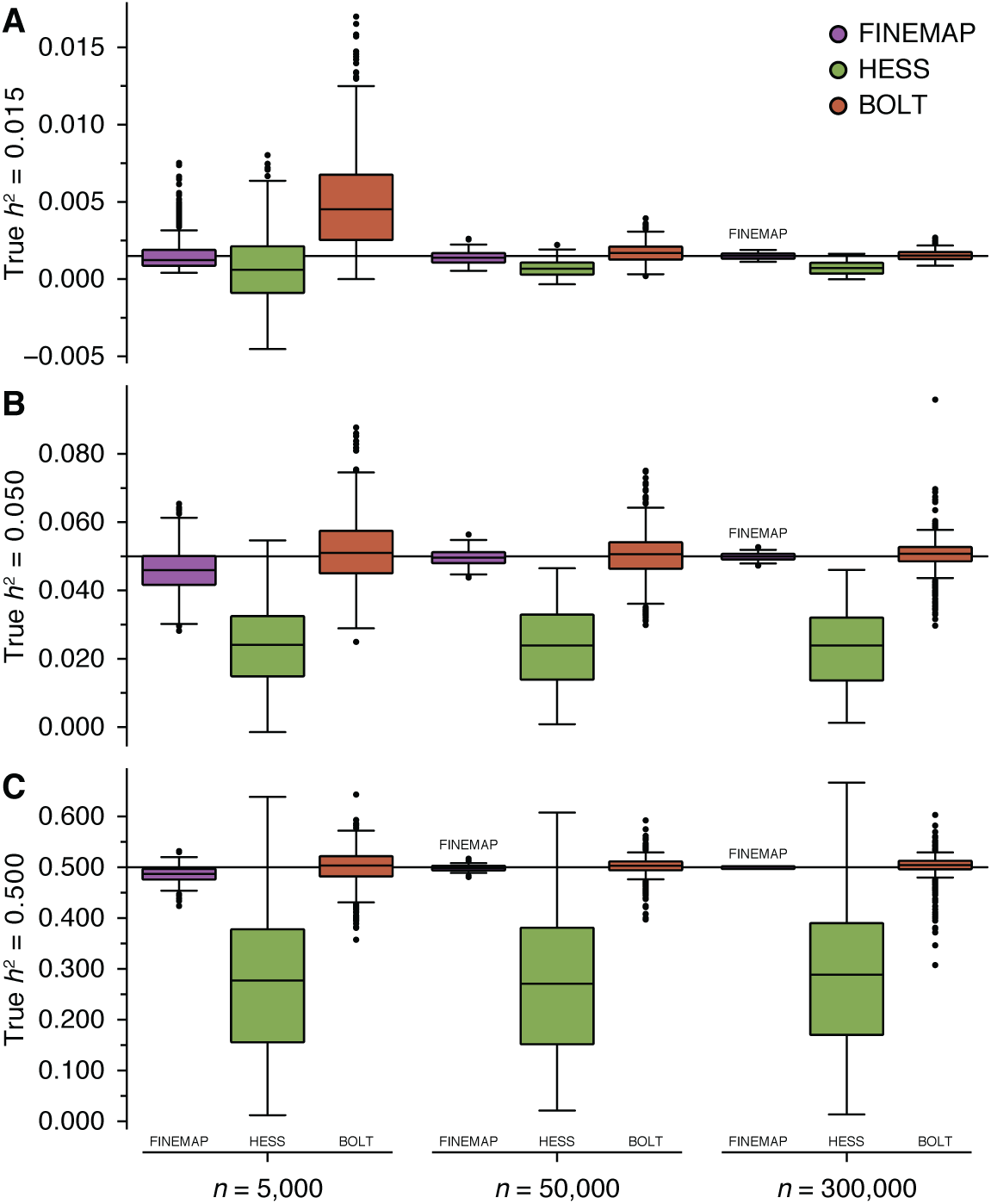
Heritability estimation on simulated data. Genotype data from UKBB over 98 GWAS regions were used for phenotype generation. Three GWAS sample sizes (*n*) and three heritability values (*h*^2^) were considered and for each combination of region, sample size and heritability, 10 datasets were generated. Each dataset included three causal variants with joint effect sizes chosen so that the three variants together account for the regional heritability in proportions of 61.8%, 25.8% and 12.4%. For each of the three estimation methods, we show the boxplots of point estimates over the datasets with true regional heritability of 0.15% (A), 5% (B) and 50% (C).

BOLT was unbiased for larger *h*^2^ (5% and 50%, Figure 1B and 1C) but strongly upwardly biased for the smallest *h*^2^ of 0.15% when *n* = 5,000 (Figure 1A). This may be because it constrains the REML-estimates to reside in the interval [0,1] and hence overestimates small heritability values. RMSE of BOLT improved with increasing sample size except for the largest heritability value, which may indicate that extreme violation of the polygenicity assumption cannot anymore be compensated with increasing information provided by larger sample size. Compared to FINEMAP, BOLT was less biased with *n* = 5,000 and larger heritability (5% and 50%) but still always had larger RMSE than FINEMAP. This suggests that the polygenic assumption of BOLT leads to less precise estimates compared to FINEMAP in cases where heritability can be attributed to a few causal variants. For *n* = 5,000, BOLT gave on average 243% and 12% higher estimates than FINEMAP in datasets with true *h*^2^ equalling to 0.15% and 5%, respectively.

HESS with regularization indicated a systematic downward bias without improvement at larger sample sizes (Figure 1). This may indicate that the regularization method applied by HESS can cause considerable underestimation of heritability in regions with only a few causal variants. To better understand this phenomenon we also ran HESS without regularization. We observed that HESS was approximately unbiased for smaller *h*^2^ (0.15% and 5%) for all sample sizes, whereas it was strongly upwardly biased at the largest *h*^2^ of 50% without improvement at large sample sizes (Figure S1 and Table S3).

Tables S2A and S2B show that FINEMAP and BOLT provided reasonably accurate uncertainty quantification of heritability estimates in the sense that the standard deviation of the estimates is close to the average standard error (BOLT) or average posterior standard deviation (FINEMAP). The coverage of the 95% intervals was good (always above 92%) for BOLT. FINEMAP achieved a similarly good coverage for larger sample sizes whereas at the smallest sample size of *n* = 5,000 and with the highest *h*^2^ of 50%, the 95% intervals of FINEMAP achieved a coverage of 87%. This discrepancy is a consequence of our priors favoring sparse models, and hence smaller heritability estimates, in case of only weakly informative data and is improved by increasing information as shown by Table S2A. For all scenarios, 95% intervals from HESS with regularization were miscalibrated because of systematic bias in heritability estimates. HESS without regularization achieved good coverage of 95% intervals for the smallest sample size of *n* = 5,000 and low *h*^2^ of 0.15% and 5%, whereas intervals for larger heritability values were miscalibrated with increasing sample sizes (Table S2C).

Figure S2A shows a comparison of PIPs between FINEMAP and CAVIARBF across 120 datasets. The strategies for handling large effect regions yield similar PIPs for both methods, but we note that CAVIARBF does not produce estimates of effect sizes or regional heritability. Figure S2B shows a comparison of PIPs from FINEMAP using different strategies to model large effect regions. FINEMAP v1.2 performs well in all settings whereas we observe many false positives when large effects are not appropriately accounted for in FINEMAP v1.1. This is because when we incorrectly assume that all effects are small, the only way to explain large regional heritability is by stipulating that also many non-causal variants contribute to the heritability.

### Biomarker and lipid-associated genomic regions in FINRISK data

GWAS on FINRISK 1997 data (*n* = 5,267) for 51 serum biomarkers and four common lipid traits identified 109 genome-wide significant regions at 63 non-overlapping loci. Using BOLT, we observed an average genome-wide heritability of 0.24 per trait ranging from 0.00 (Insulin) to 1.00 (Paraoxonase 1), whereas causal variants in trait-associated regions estimated by FINEMAP captured on average 38% of genome-wide heritability and 24% more regional heritability than the lead variant alone (Figure 2A). FINEMAP analyses further showed that the posterior expected number of causal variants is 1.77 on average, ranging from 1.11 (Interleukin 12 p70) to 5.79 (Macrophage inflammatory protein 1 beta), and that the MAF of causal variants is 0.20 on average, ranging from 0.04 (Tumor necrosis factor beta) to 0.40 (Neopterin). The region specific results are given in Table S4.

**Figure 2.**
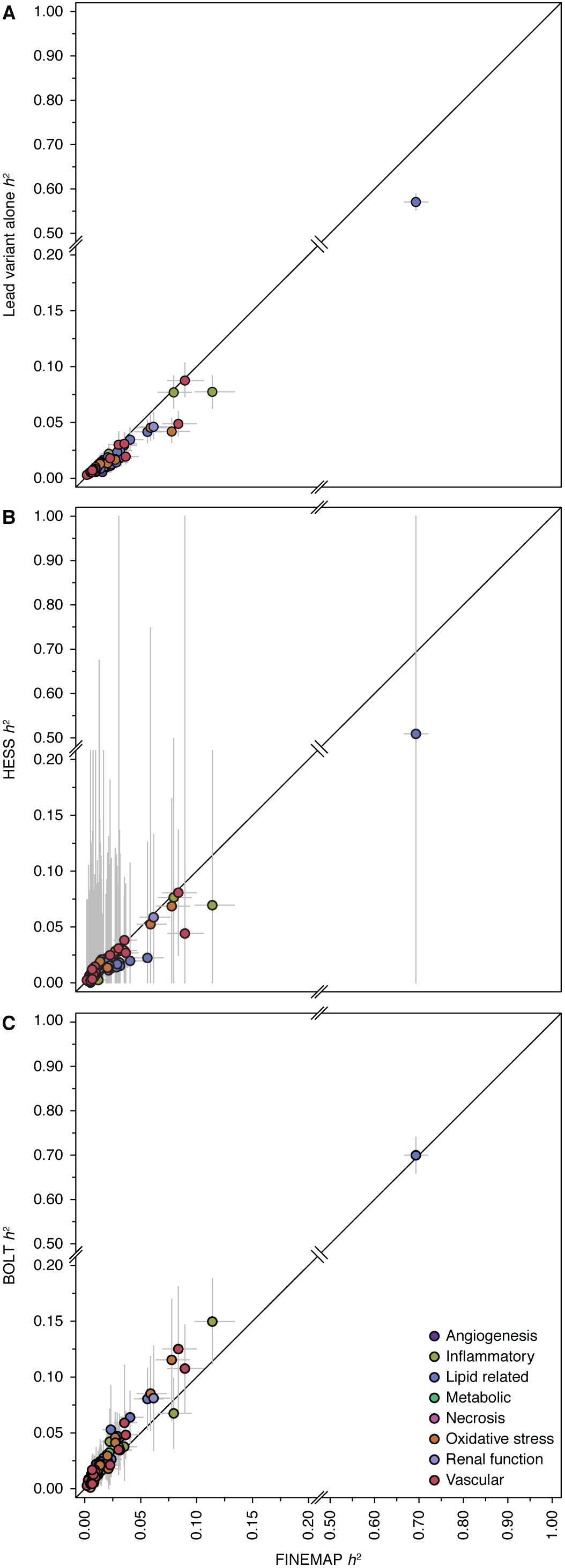
Heritability estimation on data from 51 serum biomarkers and four lipid traits. Point estimates of heritability (*h*^2^) are shown for 109 trait-associated genomic regions together with their estimated 95% intervals. Regional heritability was estimated in up to 5,167 unrelated individuals from FINRISK 1997 using variants with MAF greater than 1% and imputation quality above 0.9. Panels compare estimates from FINEMAP with the estimates from the lead variant alone (A), HESS (B) or BOLT (C).

We further evaluated whether polygenic or fixed-effects modeling implemented in BOLT and HESS, respectively, detect an excess of heritability not captured by the FINEMAP model. Heritability estimates from FINEMAP were on average 20% lower than BOLT and 40% higher than HESS with regularization (Figure 2B and 2C). BOLT and HESS provided respectively 131% and 907% wider 95% confidence intervals on average compared to 95% credible intervals from FINEMAP. Heritability estimates from HESS without regularization were systematically upwardly biased (on average over 50 times as high as FINEMAP and BOLT, Figure S3) possibly due to a relatively small sample size compared to the number of variants in the regions.

We also assessed the effect of increasing GWAS sample size on the heritability in 19 lipid-associated genomic regions. We observed that FINEMAP’s posterior expected number of causal variants grew from 1.6 using FINRISK 1997 (*n* = 5,267) to 2.42 with FINRISK 1992-2012 (*n* = 21,320). However, increasing the GWAS sample size had little effect on the captured regional heritability indicating that the newly identified causal signals account for a tiny amount of heritability and now the heritability estimates from FINEMAP were on average 23% lower than BOLT and 39% higher than HESS with regularization. At the same time, heritability estimates from BOLT and HESS were 18% and 24% lower on average with FINRISK 1992-2012 than with FINRISK 1997 alone (Figure 3).

**Figure 3.**
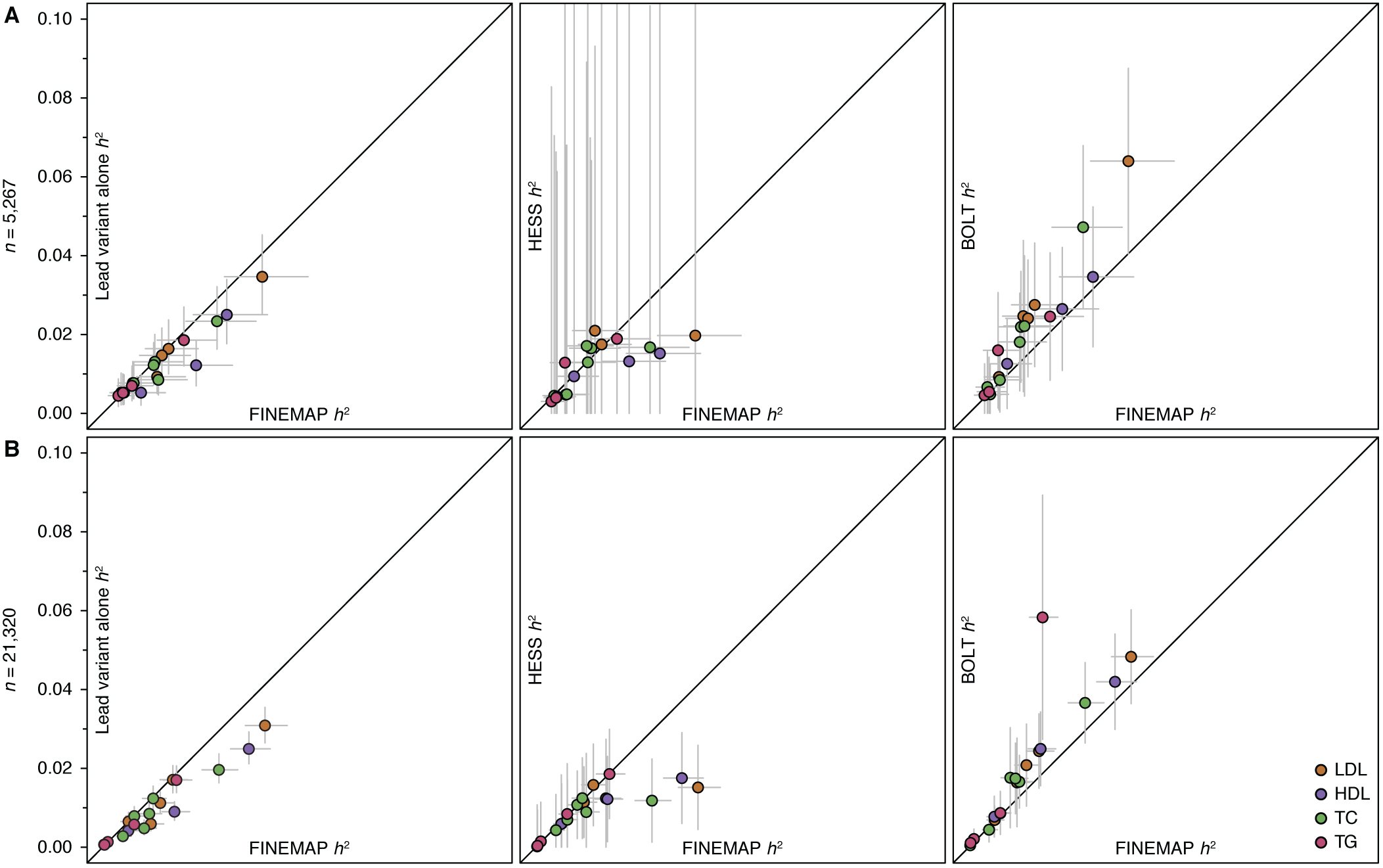
Effect of sample size in heritability estimation. Point estimates of heritability (*h*^2^) are shown for 19 lipid-associated genomic regions together with their estimated 95% intervals. Regional heritability was estimated (A) in 5,114 unrelated individuals from FINRISK 1997 and (B) in 21,320 unrelated individuals from FINRISK 1992-2012. Genomic regions included variants with MAF greater than 1% and imputation quality above 0.9. Panels compare estimates from FINEMAP with estimates from the lead variant alone (left column), HESS (middle column) or BOLT (right column).

## Discussion

Large biobanks with hundreds of thousands of samples are boosting our ability to fine-map genomic regions, that is, to probabilistically quantify which of the many correlated genetic variants have a putative causal effect on complex traits and diseases. As the power to identify causal variants and the precision of their effect size estimates increases, it is useful to be able to routinely evaluate how much of the phenotypic variance these causal variants explain together and how that compares to regional heritability estimates from models with different assumptions about the genetic architecture. To our knowledge, an estimate of the regional heritability of the fine-mapped variants is not given by existing probabilistic fine-mapping methods. In this work, we extended the FINEMAP software to output estimates of effect sizes and regional heritability by taking into account the LD structure among the variants and the posterior probability of each putative causal configuration of variants.

We performed simulations to verify our implementation and to assess how existing heritability estimation methods BOLT and HESS work in the case where a genomic region truly contains only a few causal variants. Previously, BOLT and HESS have been tested under highly polygenic architectures and here we assessed how they perform when there is considerable regional heritability but its source is not polygenic. We see this as an important piece of information since our goal is to interpret what the possible differences in the regional heritability estimates from FINEMAP, BOLT and HESS reveal about the genetic architecture of the region. On the other hand, in our simulations, we did not consider highly polygenic settings where genetic effects would be distributed among numerous variants with tiny effects. In such settings, reduced statistical power to detect individual causal variants makes fine-mapping difficult and consequently also keeps the regional heritability estimate of FINEMAP low. We expect that some existing methods, such as BOLT and HESS, could still provide reasonable heritability in such highly polygenic regions.

Our simulation results show that both FINEMAP and BOLT are nearly unbiased for larger sample sizes and provide accurate uncertainty quantification of the regional heritability estimates when there are only a few causal variants in the region. We emphasize that BOLT is derived under polygenic assumptions and therefore its application to our simulation setting may seem a potential misuse of the software, but it performed well also in our simulation setting. There was an upward bias for the smallest heritability value possibly because BOLT constrains the REML-estimates to be positive. BOLT had wider confidence intervals compared to FINEMAP estimates, likely due to our simulation setup which differed from the BOLT model and was similar to the FINEMAP model.

We also observed that the default regularization method applied by HESS induces a systematic downward bias and low precision in the estimates. The default regularization corresponds to using only the first 50 principal components of the correlation structure of the variants to remove noise present in the trailing components, especially when the LD structure does not match the GWAS summary statistics. A possible explanation for the downward bias is that the weight of the variants contributing to heritability may be weak on the leading principal components and therefore using the default settings in HESS misses important information. In our simulations, we had a perfect match between the LD structure and GWAS summary statistics and we observed that by deactivating the regularization method we could remove the bias for the smallest heritability levels. However, this could not solve the problem for the largest heritability level and led to larger confidence intervals that increased with GWAS sample size. In the biomarker data analysis, we also observed that deactivating regularization of HESS does not always work even when we have access to the original LD structure.

Our expectation in the analysis of biomarker data was that the FINEMAP heritability estimates would be bounded from below by the contribution from the variant with the lowest P-value alone and from above by polygenic and fixed-effect models, such as BOLT and HESS, that conceptually consider all variants causal. We indeed observed such pattern with FINEMAP explaining 24% more regional heritability on average than the variant with the lowest P-value alone and 20% less than BOLT. In our simulation studies BOLT overestimated the smallest regional heritability and for *n* = 5,000 BOLT gave on average 243% and 12% higher estimates than FINEMAP for heritability values of 0.15% and 5%, respectively. Therefore, a part of the higher point estimates of BOLT compared to FINEMAP in the biomarker analysis may result from BOLT overestimating and/or FINEMAP underestimating the regional heritability. Another possible explanation is that these trait-associated regions may contain a heritability contribution that we do not currently capture by fine-mapping with our sample size of about 5,000 individuals. Similar to our simulations, estimates from FINEMAP were on average 40% higher than HESS for the biomarkers. However, this time the deactivation of the regularization method of HESS did not resolve the downward bias in HESS estimates. A possible explanation is the presence of (nearly) perfectly correlated variants in the trait-associated regions making the correlation matrices not invertible or unstable in genomic regions with (nearly) perfectly correlated variants.

An ultimate goal in genetics research is to narrow down the polygenic regional heritability into individual variants contributing to heritability and to obtain accurate effect sizes for them. The extensions for effect size and heritability estimation that we have introduced in the FINEMAP software provide a computationally efficient framework to deduce a variant-level picture of the regional genetic architecture. We expect that these new features prove useful as the accuracy of fine-mapping constantly increases with rapidly growing sample size of biobank-scale studies.

## Author contributions

C.B. and M.P. designed the study. C.B. developed the software tools and conducted the analyses. A.S.H. and V.S. provided materials. S.R. and M.P. supervised the research. C.B. and M.P. wrote the manuscript. All authors reviewed the manuscript.

## Acknowledgments

We thank the participants of the FINRISK cohort and its funders: the National Institute for Health and Welfare, the Academy of Finland (139635 to V.S.), and the Finnish Foundation for Cardiovascular Research. This research was conducted with the UK Biobank Resource under application no. 22627. This work was financially supported by the Doctoral Programme in Population Health (C.B.), the Academy of Finland (288509, 294050, 312076 to M.P.; 251217 and 255847 to S.R.) and by the Academy of Finland Center of Excellence in Complex Disease Genetics. S.R. was further supported by EU FP7 project ENGAGE (201413), BioSHaRE (261433), the Finnish Foundation for Cardiovascular Research, Biocentrum Helsinki, and the Sigrid Jusélius Foundation.

## Web Resources

- BOLT, https://data.broadinstitute.org/alkesgroup/BOLT-LMM
- CAVIARBF, https://bitbucket.org/Wenan/caviarbf
- FINEMAP, http://www.finemap.me
- HESS, https://github.com/huwenboshi/hess
- IMPUTE2, http://mathgen.stats.ox.ac.uk/impute/impute_v2.html
- PLINK, https://www.cog-genomics.org/plink
- SNPTEST2, http://mathgen.stats.ox.ac.uk/genetics_software/snptest/snptest.html
- UKBB imputation, http://www.ukbiobank.co.uk/wp-content/uploads/2014/04/imputation_documentation_May2015.pdf

## References

1. Visscher, P.M. et al. 10 Years of GWAS Discovery: Biology, Function, and Translation. Am J Hum Genet 101, 5-22 (2017).

2. Manolio, T.A. et al. Finding the missing heritability of complex diseases. Nature 461, 747-53 (2009).

3. Spain, S.L. & Barrett, J.C. Strategies for fine-mapping complex traits. Hum Mol Genet 24, R111-9 (2015).

4. Yang, J. et al. Common SNPs explain a large proportion of the heritability for human height. Nat Genet 42, 565-9 (2010).

5. Loh, P.R. et al. Efficient Bayesian mixed-model analysis increases association power in large cohorts. Nat Genet 47, 284-90 (2015).

6. Bulik-Sullivan, B.K. et al. LD Score regression distinguishes confounding from polygenicity in genome-wide association studies. Nat Genet 47, 291-5 (2015).

7. Kang, H.M. et al. Variance component model to account for sample structure in genome-wide association studies. Nat Genet 42, 348-54 (2010).

8. Yang, J., Lee, S.H., Goddard, M.E. & Visscher, P.M. GCTA: a tool for genome-wide complex trait analysis. Am J Hum Genet 88, 76-82 (2011).

9. Zhou, X. & Stephens, M. Genome-wide efficient mixed-model analysis for association studies. Nat Genet 44, 821-4 (2012).

10. Pirinen, M., Donnelly, P. & Spencer, C.C.A. Efficient Computation with a Linear Mixed Model on Large-Scale Data Sets with Applications to Genetic Studies. Annals of Applied Statistics 7, 369-390 (2013).

11. Palla, L. & Dudbridge, F. A Fast Method that Uses Polygenic Scores to Estimate the Variance Explained by Genome-wide Marker Panels and the Proportion of Variants Affecting a Trait. Am J Hum Genet 97, 250-9 (2015).

12. Yang, J. et al. Genetic variance estimation with imputed variants finds negligible missing heritability for human height and body mass index. Nat Genet 47, 1114-20 (2015).

13. Finucane, H.K. et al. Partitioning heritability by functional annotation using genome-wide association summary statistics. Nat Genet 47, 1228-35 (2015).

14. Shi, H., Kichaev, G. & Pasaniuc, B. Contrasting the Genetic Architecture of 30 Complex Traits from Summary Association Data. Am J Hum Genet 99, 139-53 (2016).

15. Hormozdiari, F., Kostem, E., Kang, E.Y., Pasaniuc, B. & Eskin, E. Identifying causal variants at loci with multiple signals of association. Genetics 198, 497-508 (2014).

16. Kichaev, G. et al. Integrating functional data to prioritize causal variants in statistical fine-mapping studies. PLoS Genet 10, e1004722 (2014).

17. Chen, W. et al. Fine Mapping Causal Variants with an Approximate Bayesian Method Using Marginal Test Statistics. Genetics 200, 719-36 (2015).

18. Benner, C. et al. FINEMAP: efficient variable selection using summary data from genome-wide association studies. Bioinformatics 32, 1493-501 (2016).

19. Newcombe, P.J., Conti, D.V. & Richardson, S. JAM: A Scalable Bayesian Framework for Joint Analysis of Marginal SNP Effects. Genet Epidemiol 40, 188-201 (2016).

20. Gusev, A. et al. Quantifying missing heritability at known GWAS loci. PLoS Genet 9, e1003993 (2013).

21. Loh, P.R. et al. Contrasting genetic architectures of schizophrenia and other complex diseases using fast variance-components analysis. Nat Genet 47, 1385-92 (2015).

22. O’Hara, R.B. & Sillanpaa, M.J. A review of Bayesian variable selection methods: what, how and which. Bayesian Anal. 4, 85-117 (2009).

23. Benner, C. et al. Prospects of Fine-Mapping Trait-Associated Genomic Regions by Using Summary Statistics from Genome-wide Association Studies. Am J Hum Genet 101, 539-551 (2017).

24. Chen, W., McDonnell, S.K., Thibodeau, S.N., Tillmans, L.S. & Schaid, D.J. Incorporating Functional Annotations for Fine-Mapping Causal Variants in a Bayesian Framework Using Summary Statistics. Genetics 204, 933-958 (2016).

25. Hans, C., Dobra, A. & West, M. Shotgun Stochastic Search for “Large p” Regression. Journal of the American Statistical Association 102, 507-516 (2007).

26. Kass, R.E. & Raftery, A.E. Bayes Factors. Journal of the American Statistical Association 90, 773-795 (1995).

27. Borodulin, K. et al. Forty-year trends in cardiovascular risk factors in Finland. Eur J Public Health 25, 539-46 (2015).

28. Sudlow, C. et al. UK biobank: an open access resource for identifying the causes of a wide range of complex diseases of middle and old age. PLoS Med 12, e1001779 (2015).

29. Manichaikul, A. et al. Robust relationship inference in genome-wide association studies. Bioinformatics 26, 2867-73 (2010).

30. Surakka, I. et al. The impact of low-frequency and rare variants on lipid levels. Nat Genet 47, 589-97 (2015).

31. Blankenberg, S. et al. Contribution of 30 biomarkers to 10-year cardiovascular risk estimation in 2 population cohorts: the MONICA, risk, genetics, archiving, and monograph (MORGAM) biomarker project. Circulation 121, 2388-97 (2010).

32. Nikpay, M. et al. A comprehensive 1,000 Genomes-based genome-wide association meta-analysis of coronary artery disease. Nat Genet 47, 1121-30 (2015).

33. Jostins, L. et al. Host-microbe interactions have shaped the genetic architecture of inflammatory bowel disease. Nature 491, 119-24 (2012).

34. Willer, C.J. et al. Discovery and refinement of loci associated with lipid levels. Nat Genet 45, 1274-83 (2013).

35. Schizophrenia Working Group of the Psychiatric Genomics Consortium. Biological insights from 108 schizophrenia-associated genetic loci. Nature 511, 421-7 (2014).

36. Morris, A.P. et al. Large-scale association analysis provides insights into the genetic architecture and pathophysiology of type 2 diabetes. Nat Genet 44, 981-90 (2012).

37. Locke, A.E. et al. Genetic studies of body mass index yield new insights for obesity biology. Nature 518, 197-206 (2015).

38. Tang, W.H. et al. Clinical and genetic association of serum paraoxonase and arylesterase activities with cardiovascular risk. Arterioscler Thromb Vasc Biol 32, 2803-12 (2012).

